# Genome Report: Improved chromosome-level genome assembly of the American cockroach, *Periplaneta americana*

**DOI:** 10.1101/2025.03.14.643325

**Authors:** Rachel L. Dockman, Tyler J. Simmonds, Kevin J. Vogel, Scott M. Geib, Elizabeth A. Ottesen

## Abstract

The American cockroach, *Periplaneta americana*, is a cosmopolitan insect notorious for thriving among humans undeterred by attempts to eliminate it. The traits that contribute to its ubiquity as an opportunistic pest, such as long lifespan, expansive neurosensory capacity, and nutritional flexibility, also make *P. americana* an excellent invertebrate model organism with a long history in neuroscience and physiological research. Current genetic resources available for *P. americana* highlight its large, complex genome and richly diverse transcriptional capabilities, but fall short of producing a complete, chromosome-level genome. Here, we present a high-quality *de novo* genome assembly of a laboratory-raised adult female *P. americana* using a combination of high fidelity PacBio long reads and Hi-C sequencing. The final 3.23 Gb genome was assembled with chromosomal resolution into 17 scaffolds, consistent with previous karyotype analysis, and has a scaffold N50 of 188.1 Mb and genome BUSCO score of 99.7%. This assembly includes a chromosome that was missing from the previous reference genome for this species. Protein prediction and annotation were performed via the NCBI Eukaryotic Genome Annotation Pipeline, which identified 16,780 protein-coding genes and generated an annotation BUSCO score of 97.8%. Ortholog comparisons with available Blattodea assemblies highlight the expanded chemosensory and immune capabilities of *P. americana* compared to termite relatives. This genome assembly is a valuable tool for facilitating future research on the biology and evolution of this remarkable insect.

## 2 Introduction

The American cockroach (*Periplaneta americana*) is a notorious pest found across the world, living and thriving alongside humans in widely variable environmental conditions. Despite their unsavory reputation, cockroaches are of considerable interest to researchers across disciplines. Cockroaches combine the large size, complex physiology, and tractable nature of mice with the simpler and cost-effective care requirements characteristic of model insects.

There is a rich body of work compiled throughout the last century describing basic cockroach biology, covering topics ranging from external morphology to internal physiology [1–4]. Early research leveraging *P. americana* as a model system has enhanced understanding of nervous system function, connectivity, and regeneration [5–9] and discovered unique traits that allow *P. americana* to survive extreme environmental stressors, such as endosymbiont-mediated nitrogen cycling [10, 11]. Studies on the cockroach immune system have linked its expansive repertoire of immune-associated proteins to common allergens [12, 13] and mechanisms of pesticide resistance [14–16]. The extensive antimicrobial and regenerative capabilities of *P. americana* have also earned it respect in both ancient and modern Chinese tradition as an important medicinal insect [17, 18]. Modern sequencing technologies and genetic techniques have further supported the use of *P. americana* as a model organism. Its susceptibility and robust response to RNA inhibition (RNAi) via multiple administration methods, *P. americana* is an especially useful organism for deciphering gene involvement in insect physiological development and pesticide resistance, among other investigations [19–21]. *P. americana* also shows potential as an emerging insect model for host-microbiome interactions. The gut microbiome of omnivorous cockroaches reflects that of humans and omnivorous mammalian model systems [22–25], and germ-free nymphs can be easily produced via ootheca sterilization, a highly desirable trait for defined-community research interests [26–29].Altogether, these traits support the research potential of this insect and argue towards the necessity of a complete, high-quality American cockroach genome. While there are two previous genome assemblies publicly available, the assembly presented by [30] lacks publicly available protein annotations and is limited by the short-read technology available at the time and the assembly prepared by [13], while a significant improvement, contains more scaffolds than supported by karyotype analysis (male/XO: 33 diploid; female/XX: 34 diploid) [31].

Here, we present the first chromosome-level assembled genome of *P. americana*. We used long-read high fidelity (HiFi) PacBio sequencing in addition to chromatin contact mapping (Hi-C) to produce a genome scaffolded into a 17-chromosome assembly, consistent with previous karyotype findings [31]. This high-quality genome assembly is an important tool for facilitating future genetic research in the American cockroach.

## 3 Materials and Methods

### 3.1 Insect origin and selection

An adult female *Periplaneta americana* individual was selected from a stock colony maintained at the University of Georgia by the Ottesen laboratory; this colony has been maintained the laboratory for 10 years and originated in another long-term laboratory colony of unknown origin. The specimen was flash frozen in liquid nitrogen and shipped on dry ice to the United States Department of Agriculture - Agricultural Research Service (USDA-ARS) – Pacific Basin Agricultural Research Center (PBARC) in Hilo, Hawaii.

### 3.2 Sample preparation and sequencing methods

For PacBio sequencing, high molecular weight (HMW) DNA extraction was performed from insect leg tissue using the Qiagen MagAttract HMW DNA Kit (Qiagen, Hilden Germany). DNA was sheared with the Diagenode Megaruptor 2 (Denville, New Jersey, USA) then prepared for PacBio sequencing using the SMRTBell Express Template kit 2.0 (Pacific Biosciences, Menlo Park, California, USA). The library was size-selected prior to sequencing on a Sequel II System (Pacific Biosciences, Menlo Park, California, USA) using Binding Kit v2.0, Sequencing kit v2.0, and SMRT Cell 8M. To target HiFi reads, the library was sequenced using a 30-hour movie time on three SMRTcells. Raw subreads were converted to HiFi data by processing with CCS to call a single high quality consensus sequence for each molecule, using a 99.5% consensus accuracy cutoff.

For Hi-C sequencing, the Arima Hi-C kit (Arima Genomics, San Diego, California, USA) was used to crosslink leg tissue DNA and perform proximity ligation, following the Arima Hi-C low input protocol. After proximity ligation, DNA was sheared with a Diagenode Bioruptor then size-selected for 200-600bp DNA fragments. The Swift Accel NGS 2S Plus kit (Integrated DNA Technologies, Coralville, Iowa, USA) was used to prepare an Illumina library from the size-selected DNA, which was then sequenced with an Illumina NovaSeq 6000 (Illumina, San Diego, California, USA).

### 3.3 RNA-seq

Transcriptomic data was obtained for the midgut, hindgut, and fat body of 10 individual cockroaches for a dietary experiment investigating the impact of carbohydrate source on the microbial metatranscriptome and host transcriptome (BioProject: PRJNA1105088). Data were obtained as 150bp paired end reads on Illumina NovaSeq from Novogene Corporation in Sacramento, California. The Joint Genome Institute (JGI) programs BBduk and BBSplit (https://jgi.doe.gov/data-and-tools/software-tools/bbtools/) and were used to remove sequencing adapters and screen for initial *Blattabacterium* contamination, and SortMeRNA was used to remove ribosomal RNA reads [32]. Cleaned RNA reads were aligned to the unfiltered genome for contaminant filtering.

### 3.4 Genome assembly and polishing

The first genome assembly was generated using hifiasm v0.19.6-r595 Hi-C integration to obtain primary, alternate, and haplotype-phased assemblies [33]. Three flow cells of PacBio HiFi data were obtained from long-read sequencing and concatenated into a single fastq file for assembly and used with Hi-C data obtained from the same source insect. The primary assembly was selected for polishing and contigs were filtered to retain those with coverage reported from hifiasm as between 6X and 30X.

Contamination filtering was performed as described by Lu and Salzberg [34]. Contigs were separated into individual files, then fragmented with SeqKit v2.5.1 into 100 bp pseudo-reads with 50 bp overlaps [35]. These pseudo-reads were fed through Kraken v2.1.3 with default parameters to align to the default eukaryotic and prokaryotic databases, and contigs identified as *Blattabacterium* or with 70% identity assigned to *Homo sapiens* were discarded [36]. Remaining contigs were masked using RepeatMasker v4.1.5 (https://www.repeatmasker.org/RepeatMasker/) with Dfam library v3.7 prior to RNA-seq alignment with HISAT2, and contigs with average read depths exceeding 10,000 were removed [37, 38].

The Arima Genomics mapping pipeline (https://github.com/ArimaGenomics/mapping_pipeline) was used to map Hi-C data to the filtered assembly. The pipeline described utilizes the programs BWA-MEM for separate alignment of the paired Illumina Hi-C reads, the Picard (https://broadinstitute.github.io/picard/) “MarkDuplicates” command for PCR duplicate removal, and SAMtools for file sorting and handling [39, 40]. The resulting Hi-C alignment files and the hifiasm assembly were fed into Yet Another Hi-C Scaffolding tool (YaHS) without breaking contigs, and the resulting contact map was visualized with Juicebox for manual curation [41, 42]. The corrected genome was exported and screened for possible telomeres using tidk (https://github.com/tolkit/telomeric-identifier) and repeat regions identified *de novo* (AACCTAACCT) were graphed [43]. An additional round of polishing was performed in Juicebox to correct repeat-heavy telomeric loci, and completeness of the final scaffolded genome was assessed with the Insecta set (version odb10) of Benchmarking Universal Single-Copy Orthologs (BUSCOs) [44].

### 3.5 Mitochondrion identification

The program MitoHiFi v2 was used to identify a consensus mitochondrial genome sequence from the original hifiasm assembly [45]. The NCBI reference sequence NC_016956.1 was selected as the reference *P. americana* mitochondrion for identification [46].

### 3.6 Repeat modeling and masking

For in-depth repeat identification, RepeatModeler v2.0.4 was used to create an *ab initio* repeat library specific to *P. americana,* which was then separated into libraries of known and unknown repeat families [47]. Using the script repclassifier.sh (https://github.com/darencard/GenomeAnnotation/blob/master/repclassifier), unknown repeats were iteratively re-annotated for seven rounds, when the percent of repeats classified as “known” rather than “unknown” plateaued. Four rounds of repeat masking using RepeatMasker v4.1.5 were then performed as described in (https://darencard.net/blog/2022-07-09-genome-repeat-annotation/), during which simple repeats were identified and masked first, followed by insect-specific Dfam repeats, known *P. americana* repeats, and lastly unknown *P. americana* repeats. The output from the four RepeatMasker rounds were combined to generate an overall masked genome, annotation files, and table describing the repeats, and repeat landscapes were summarized with the script parseRM.pl (https://github.com/4ureliek/Parsing-RepeatMasker-Outputs).

### 3.7 Genome annotation

The finished assembly was submitted to NCBI for structural and functional gene annotation via the automated Eukaryotic Genome Annotation Pipeline v10.3. Evidence fed into the GNOMON gene prediction tool included existing RNA-seq data and transcriptome assemblies for *P. americana*, NCBI RefSeq protein sets from *Acromyrmex echinatior*, *Hyalella azteca, Acyrthosiphon pisum, Caenorhabditis elegans, Tribolium castaneum, Drosophila melanogaster*, and *Apis mellifera*, as well as Insecta and *P. americana* GenBank protein sets. Details of the annotation release (GCF_040183065.1-RS_2024_10) are available at https://www.ncbi.nlm.nih.gov/refseq/annotation_euk/Periplaneta_americana/GCF_040183065.1-RS_2024_10/.

### 3.8 Blattodea comparison

The annotations obtained for this assembly were compared with Blattodea annotations available on NCBI for a previous *P. americana* genome (GCA_025594305.2) [13], *Zootermopsis nevadensis* (GCF_000696155.1) [48], *Blattella germanica* (GCA_003018175.1) and *Cryptotermes secundus* (GCF_002891405.2) [49], *Coptotermes formosanus* (GCA_013340265.1), and *Diploptera punctata* (GCA_030220185.1) [50]. OrthoFinder v2.5.5 was used with GENESPACE v1.3.1 in R Studio to identify orthologous groups and generate riparian plots [51–53]. The distribution of orthologous gene sets between and across genomes were visualized with the R package UpSetR v1.4.0 [54]. GO terms assigned to *P. americana* genes that were sorted into orthologous gene sets were used for GO enrichment analysis in R with the package clusterProfiler v4.6.2 [55, 56].

## 4 Results/Discussion

### 4.1 Assembly

American cockroaches have XX/X0 sex chromosome systems [57], so a single female insect was selected for PacBio SMRTBell sequencing, producing over 6.6 million reads covering 90.6 Gb. Paired-end 150 bp sequences generated by Hi-C sequencing produced an additional 217.3 Gb of sequence data. These data were assembled with hifiasm in Hi-C mode to produce a primary genome assembly containing 1818 contigs with a contig N50 over 40 Mb and total length of 3.36 Gb, close to the genome size of 3.338 Gb predicted through previous flow cytometry work [58]. In addition, we identified a complete consensus mitogenome (**Figure S1**). We evaluated the assembly quality via read coverage and depth (**Figure 1**) prior to filtering and found that 96.19% of the genome assembly was contained in contigs with coverage between 6x and 30x. Subsequent contamination filtering using Kraken2 identified 19 contigs as derived from *Blattabacterium*, the cockroach-associated endosymbiont. All contigs flagged as *Blattabacterium* DNA in our genome assembly were self-contained, suggesting that these reads stemmed from contamination rather than integration into *P. americana*’s genome, and they were therefore removed. Further filtering based on RNA-seq alignment decreased the number of contigs prior to scaffolding to 243 with a N50 of 42.4 Mb, overall producing a high-quality contig-level genome with no detectable contamination.

**Figure 1:**
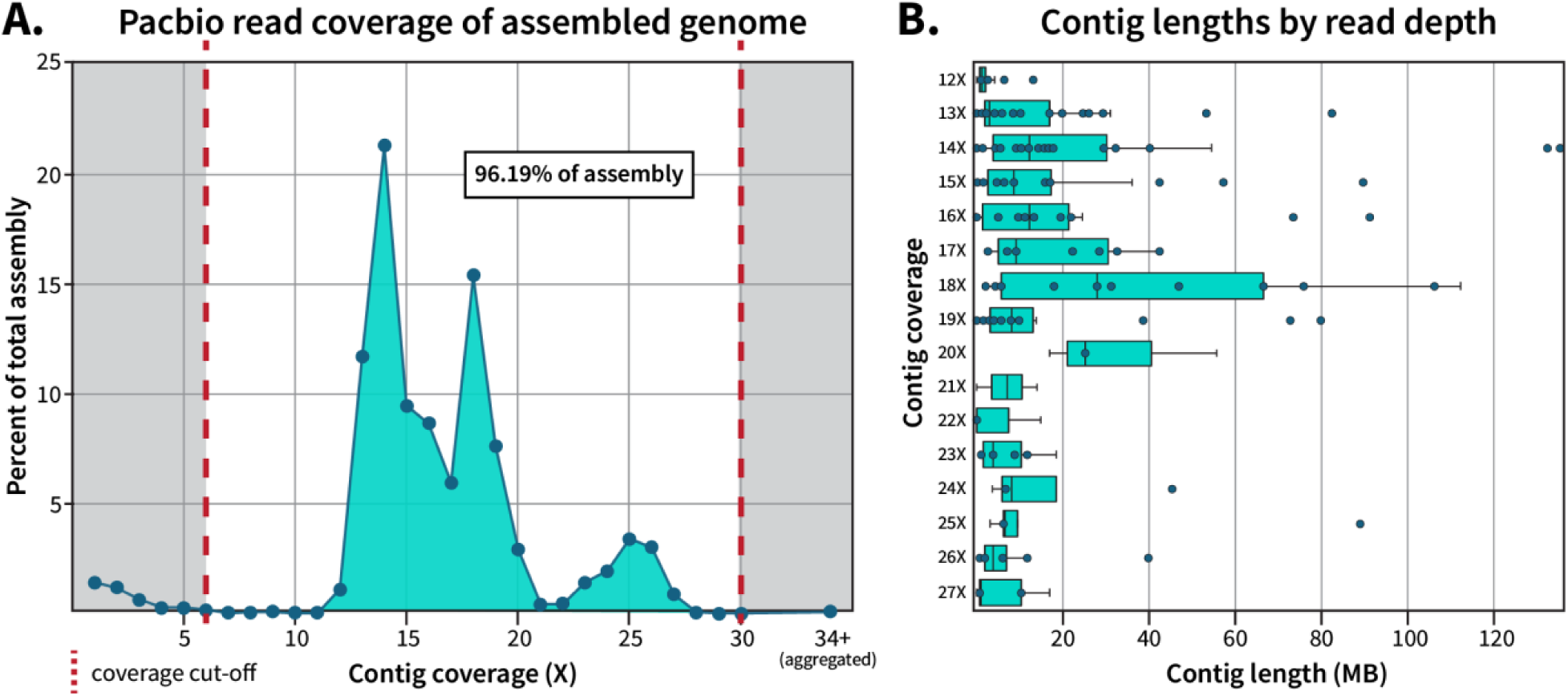
Quality profile of initial contig and final scaffold hifiasm assembly. **(A)** Pacbio reads were mapped to the primary genome assembly to determine overall coverage of individual contigs, and the percent each coverage level contributed to the total genome size was plotted. **(B)** The length of contigs per coverage level were plotted, with individual contigs represented as dots. Retained contigs with coverage between 6X-11X (n=73) or 28X-30X (n=5) were excluded from plotting in **(B)** due to short lengths.

Initial scaffolding with YaHS assigned 95.67% of the genome into 17 scaffolds (**Figure S2**), with manual curation in Juicebox increasing this percentage to 98.93% (**Figure 2A**). Telomere analysis identified at least one telomeric region, determined *de novo* as AACCTAACCT, for each of the chromosome-sized scaffolds, with 10 scaffolds flanked on both ends and 6 scaffolds flanked on one end (**Figure 2B**). Scaffold #7 contains a long sequence of probable telomeric repeats embedded within a large contig. While this may indicate an assembly error, at this time we do not have evidence to support adjusting its placement. The 74 remaining contigs were unable to be matched to just one scaffold, likely due to a high density of centromeric repeat regions and were therefore left unplaced. BUSCO analysis of the putative chromosomes found 99.7% of Insecta BUSCOs were present and complete in this genome assembly, of which only 1.5% were duplicated (**Figure S3, Table 1**).

**Figure 2:**
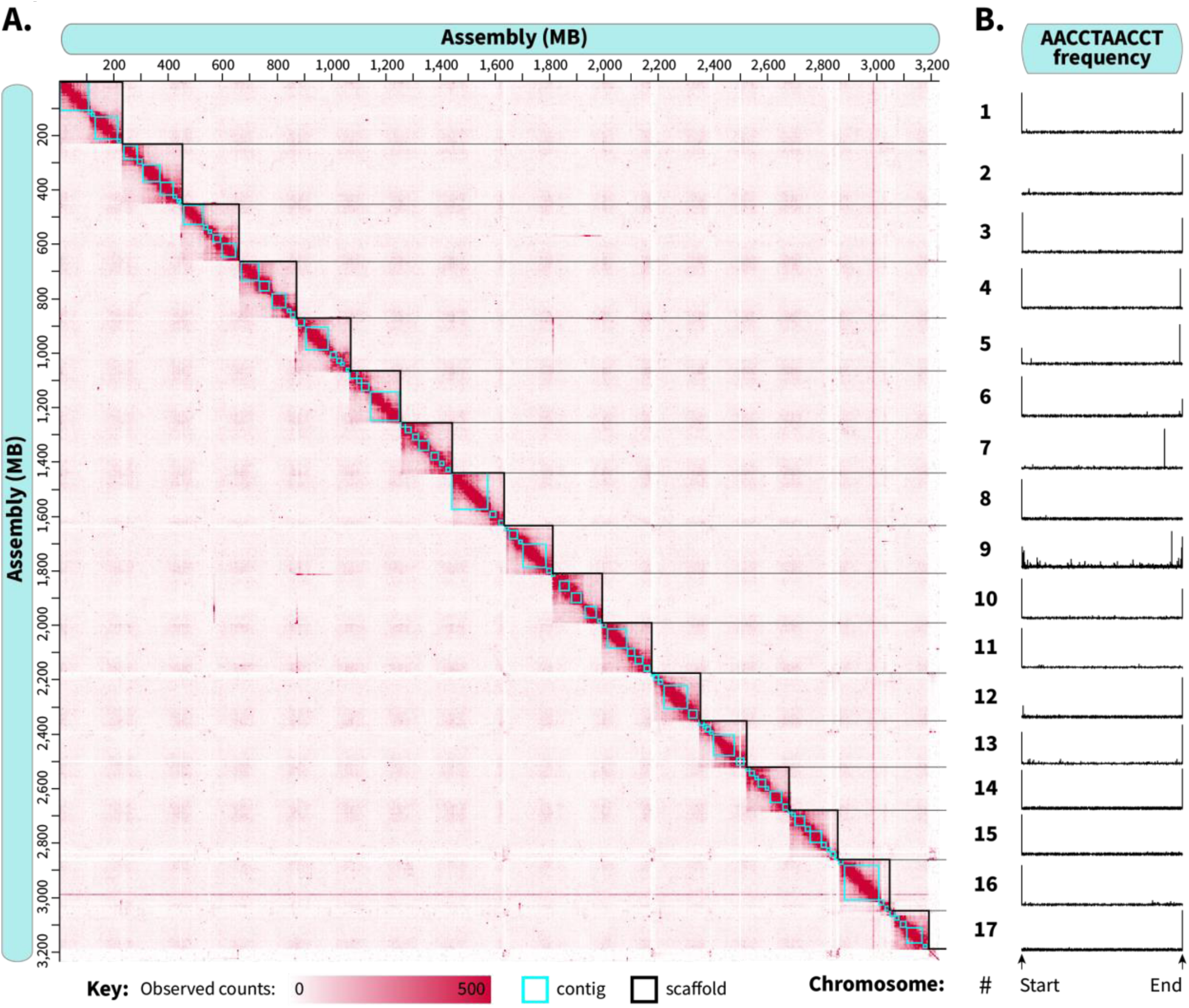
Chromatin-contact sequencing produced an assembly with near-chromosomal resolution. Following scaffolding with Yet Another Hi-C Scaffolder (YAHS), scaffold boundaries and contig placement were adjusted in Juicebox to optimize chromatin contacts for the 17 chromosome-level scaffolds. Final chromosomal boundaries are shown in the heatmap in **(A),** and the occurrence of telomeric sequences within each chromosome are displayed in **(B).**

**Table 1:**
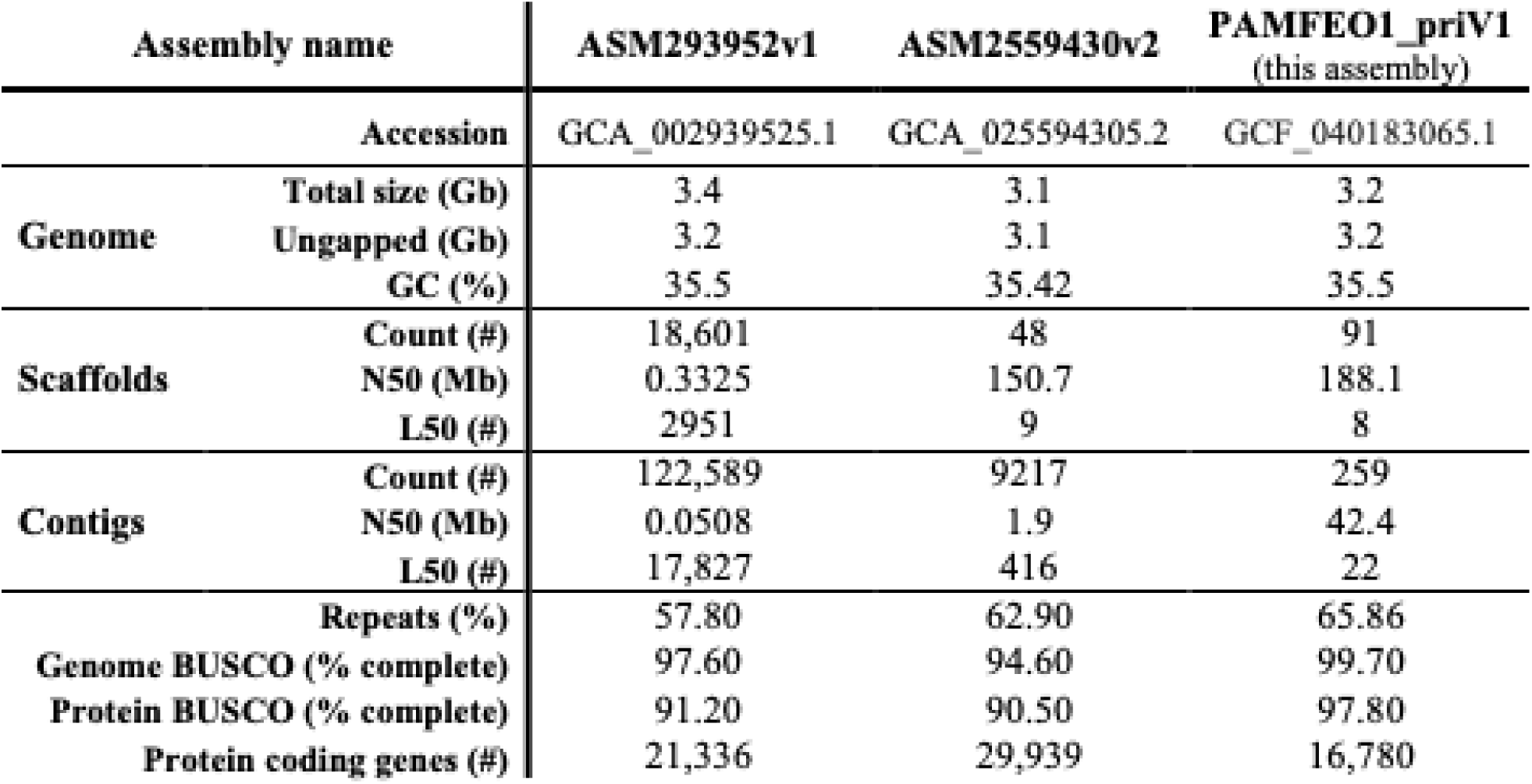
Genome assembly comparison with previous *Periplaneta americana* assemblies available on NCBI.

In summary, the assembly presented here is considered high quality across a number of standard metrics. Through a combination of long-read and chromosomal contact sequencing data, we successfully scaffolded 98.9% of this 3.23 Gb assembly into 17 chromosome-scale scaffolds, values which are supported by previous karyotype and flow cytometry findings for *P. americana* of 17 haploid (male/female: 33/34 diploid) chromosomes and a predicted genome size of 3.338 Gb [31, 57, 58]. This represents an improvement on the currently available *P. americana* genome assemblies in NCBI (**Table 1**), which resemble this assembly in length and GC content but have lower contiguity and BUSCO scores [13, 30]. Therefore, we argue that this genome assembly qualifies as both comprehensive and chromosomally resolved.

### 4.2 Repetitive DNA elements

Overall, 50.86% of the genome was identified as repetitive content classified as DNA elements, simple repeat regions, or retroelements, with an additional 15% of the genome determined to be *P. americana*-specific repeats that remain otherwise unclassified **(Figure 3A, Table S1**). The DNA transposon and retroelement subgroups contributed similarly to overall repeat content, comprising 24.3% and 20.05% respectively of the genome, but differ in their overall Kimura divergence landscapes (**Figure 3B)**. The DNA elements found in this genome primarily belong to the Tc1-Mariner and hobo-Activator-Tam3 subfamilies with higher relative abundances at a Kimura substitution level of 5% (**Figure 3C**). In contrast, the primary retrotransposon class of long interspersed nuclear elements (LINEs) contained two peaks in Kimura substitution, with CR1, L1, L2, and CRE subgroups showing elevated substitution levels around 5% while RTE-clade retrotransposons represent a more ancient repeat lineage, with substitution levels peaking between 31-32% **(Figure 3D)**. Short interspersed nuclear elements (SINEs) and retroelements containing long-terminal repeats (LTRs) were less commonly identified, only making up 3.47% and 0.94% of the genome respectively. Most SINEs contained internal promoters derived from tRNA and showed more gradual patterns of divergence, with a relatively stable plateau between 5% and 15% divergence before tapering (**Figure 3E).** Repeat content varies widely even between insects from the same family, but generally Blattodea species show similar repeat distributions with especially low LTR content and expanded LINEs, with larger genomes correlated to higher repeat content [59].

**Figure 3:**
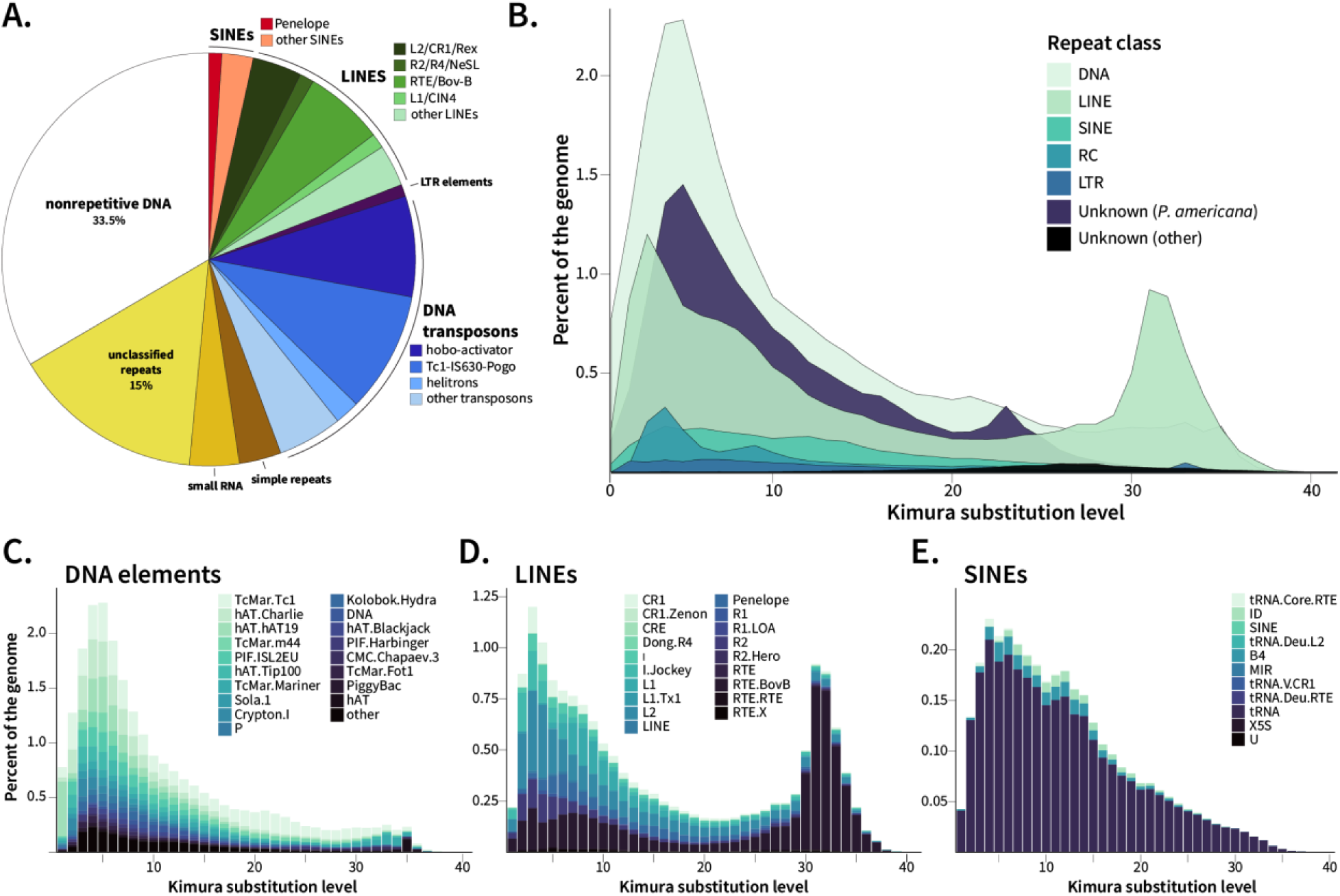
Repeat summary of *P. americana*. RepeatMasker and RepeatModeler were used to identify **(A)** the abundance of repetitive element families present in this assembly and **(B)** the relative abundance of each repeat class versus Kimura substitution level. The repeat landscapes for the classes **(C)** DNA elements, **(D)** LINEs, and **(E)** SINEs were further visualized at the repeat subtype level.

### 4.3 Orthology analysis between Blattodea species

The order Blattodea encompasses both termites and cockroaches, with over 4700 species identified in NCBI’s taxonomy repository. Despite these many representatives, only 12 species have sequenced genomes [13, 30, 48–50, 60–63], of which half have publicly available annotations uploaded to NCBI (**Figure 4A**). The 3.2 Gb genome of *P. americana* is the largest among sequenced Blattodea and more than double the size of available termite genomes (**Figure 4B**), consistent with two previous *P. americana* genome assemblies. This assembly has the highest contig N50 among these Blattodea (**Figure 4C)** and the second highest scaffold N50 behind *E. pallidus* [60] **(Figure 4D).** Compared to those genomes with available annotations, *P. americana* encodes more protein-coding genes than termites, but had less reported than the other cockroach genomes (**Figure 4E**). It is unclear whether this difference between annotations in our assembly and the other cockroach assemblies is biological or a result of annotation technique; this assembly was annotated by NCBI’s Gnomon pipeline which produced 16,780 protein-coding genes, while previous *P. americana* genomes reported 21,336 (annotations not available) and 29,939 (annotations available) protein-coding genes [13, 30].

**Figure 4:**
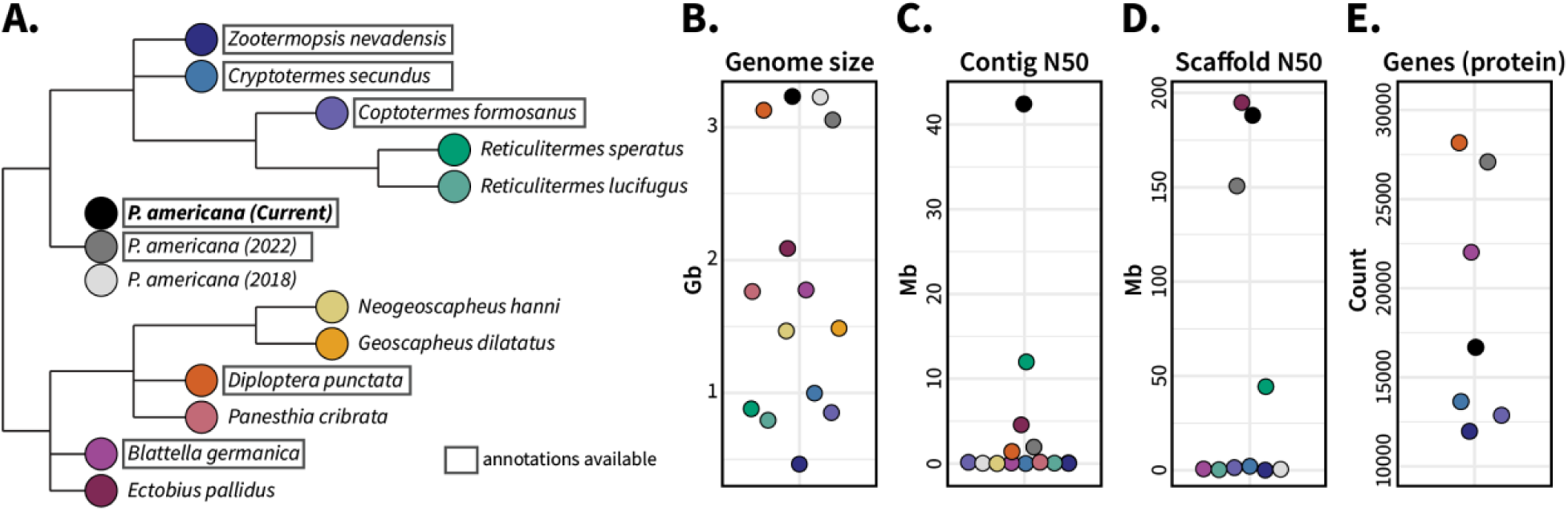
Available Blattodea genomes from NCBI. Phylogeny of sequenced Blattodea genomes as they appear in the NCBI taxonomy browser is presented in **(A)**, in addition to their **(B)** genome size, **(C)** contig N50, **(D)** scaffold N50, and (**E)** number of protein-coding genes (if available). Points on the genome statistic plots **(B-E)** correspond to the colors in **(A)** and species in (**A**) with protein annotations available on NCBI are boxed.

We chose to use the genome annotations available on NCBI for ortholog analysis, which included three members of Termitidae (*Zootermopsis nevadensis, Cryptotermes secundus, Coptotermes formosanus*), two Blaberoidea (*Blattella germanica, Diploptera punctata*) and a previous *P. americana* genome [13]. Initial OrthoFinder results included both *P. americana* assemblies, identifying 16,711 orthologous gene families among these seven annotated genomes (**Supplemental File 1**). Of these 16,711 gene families, 2,311 were shared across all seven Blattodea with an additional 2,220 gene families shared by all assemblies excluding the previous *P. americana* assembly (**Figure S4**). While both *P. americana* genomes shared 391 gene families absent in the other species, 198 gene families were only present in our assembly (**Figure S4**). We further evaluated the differences between both *P. americana* assemblies via synteny analysis (**Figure 5A**) and found that, while most chromosomes in our assembly were captured in their entirety by 1-2 scaffolds in the previous assembly, our chromosome 10 and half of chromosome 14 were missing from the other *P. americana* annotations. Proteins encoded in chromosome 10 encompass a wide range of functions with immune (Dscam, toll-like receptors, leucine-rich repeat proteins), neurologic (GABA transport, neurotrophic factors), endocrine (vitellogenin synthesis, sterol binding proteins), and nutritional (xanthine dehydrogenase, salivary peptide) importance. Additional synteny analysis between our *P. americana* genome and other cockroaches (**Figure 5B**) and termites (**Figure 5C**) supported the existence of these regions. As a result, the previous *P. americana* assembly was excluded from further cross-*Blattodea* orthogroup analysis.

**Figure 5:**
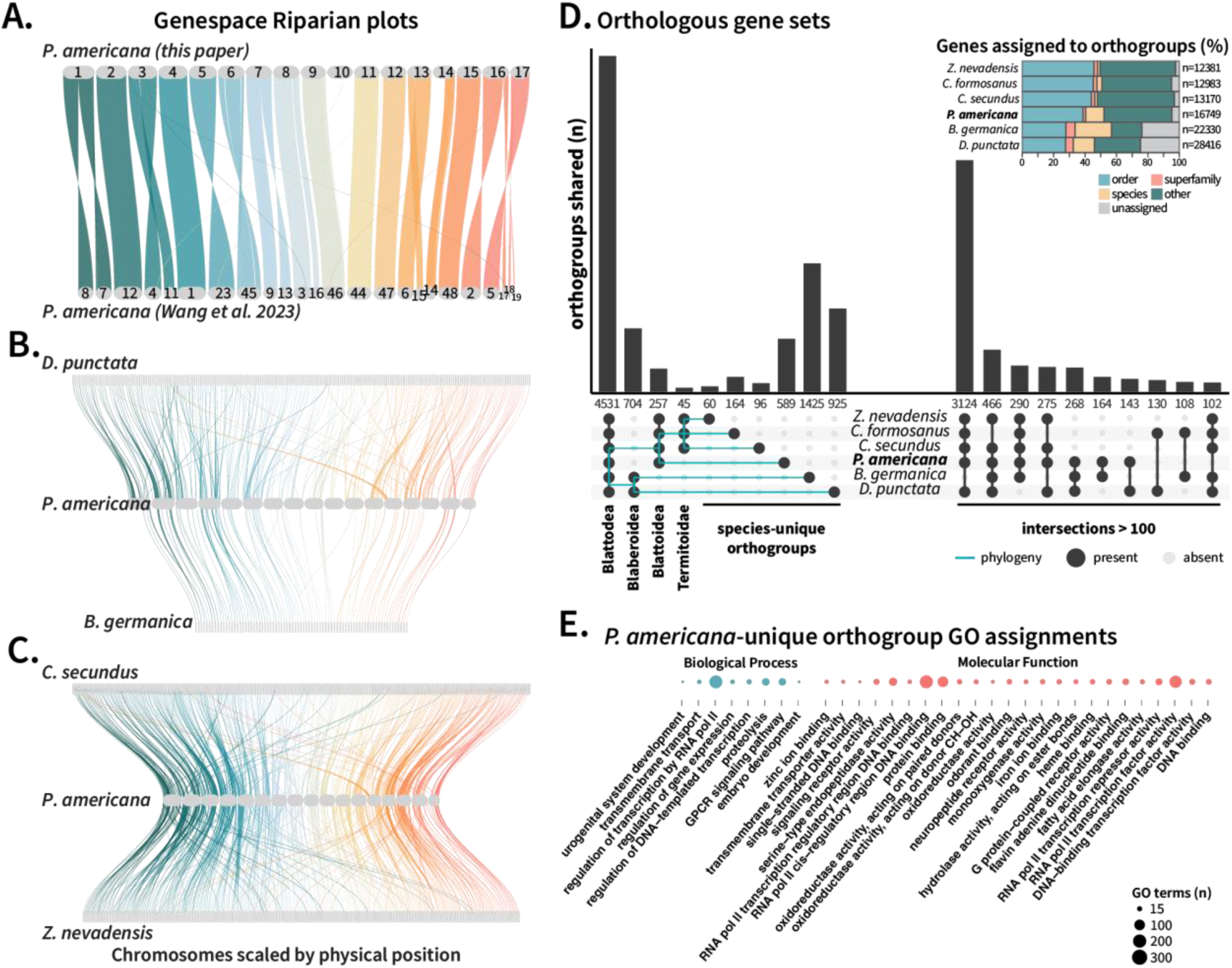
Ortholog analysis within Blattodea. Protein GFF3 files available on NCBI for Blattodea species were compared using OrthoFinder and GENESPACE. Riparian plots were generated to assess synteny between the *P. americana* genome presented here and **(A)** a previous *P. americana* assembly (GCA_025594305.2), **(B)** the cockroaches *Diploptera puntata* (GCA_030220185.1) and *Blattella germanica* (GCA_003018175.1), and **(C)** the termites *Cryptotermes secundus* (GCF_002891405.2) and *Zootermopsis nevadensis* (GCF_000696155.1). (**D**) Orthologous gene clusters shared by or unique to the analyzed genomes were visualized via UpSet plot, organized by phylogeny on the left and the next 10 largest sets on the right and summarized by (**inset D)** the percentage of genes assigned to shared or unique orthogroups. Genes belonging to the 589 orthogroups identified as *P. americana*-unique were analyzed by their associated GO terms, and (**E**) biological processes and molecular functions with at least 15 occurrences were visualized.

Recalculating OrthoFinder statistics produced 15,491 total gene families, of which 4,531 orthogroups were shared by all species (**Figure 5D**). These gene families, which are shared at the order level, comprised large fractions of each insect’s total gene count, ranging from 5,674 genes (45.8%) in *Z. nevadensis* to 7,860 genes (27.7%) in *D. punctata* **(Figure 5D inset)**. These core gene families encompass a wide range of functions necessary to life and common in insects, with GO terms relating to gene expression and genome maintenance especially prevalent in this subset (**Figure S5**). The full list of GO terms assigned to these shared gene families can be found in **Supplemental File 2**.

The percentage of genes unable to be assigned to orthogroups varied between Blaberoidea and Blattoidea (**Figure 5D inset**). The termite genomes and our *P. americana* assembly had between 95-97.7% of their genes assigned to orthogroups, while 75.3% and 76.2% of *D. punctata* and *B. germanica* genes, respectively, were assigned to orthogroups (**Figure 5D inset**). Unassigned genes are considered unique both between analyzed genomes and within a single genome, so it is unclear whether these genes stem from expanded single-copy genes within the cockroaches or are misannotated as genes without conferring any biological function. Possibly, the annotations of the termites and our *P. americana* are overly conservative, but gene finding programs rely on homology comparison that requires more sequenced and annotated Blattodea genomes for effective performance. Despite these questions, the difference between cockroaches and termites in their species-specific gene families highlight the gene expansion occurring as these two groups diverged. Our *P. americana* assembly, which encodes over 3,000 more genes than the termite assemblies, has 11.7% of its genes assigned to the 589 *P. americana*-unique gene families, a substantial increase compared to the 1.5-3.2% range in the three termite assemblies **(Figure 5D inset**). This expansion of genes within a single species also occurs in *B. germanica* and *D. punctata*, which have 23.2% and 13.5% of their genes respectively assigned to species-specific gene families. Since ortholog analysis is dependent on the data supplied, it is difficult to determine whether this difference stems from biological cockroach-termite delineation or is a consequence of poor representation of closely related cockroach species. Nonetheless, these results, in combination with the large size of this cockroach genome (**Figure 4B**) and the late spike in LINE retrotransposon divergence **(Figure 3D),** suggests that acquisition and expansion of new genes contributed to cockroach divergence.

We performed GO term analysis of the 589 orthologous gene families present in our *P. americana* assembly but absent in the other Blattodea species (**Figure 5E**). GO terms related to gene expression wand genomic maintenance were most enriched, reflecting the GO enrichment in shared gene families (**Figure S5**). Further investigation revealed that 157 of these orthogroups were associated with zinc finger family genes, likely skewing the GO analysis results and masking other expanded functions of interest. As an alternative approach, we evaluated the semantic similarity of named genes and/or GO terms (if name is missing) of these *P. americana*-unique gene families to shed light on the function of these expanded genes (**Figure S6**). These gene families were enriched in immune and digestive functions, such as lipopolysaccharide recognition, protease activity, odorant binding, and lipase activity (**Figure S6**), which may have facilitated cockroach divergence from the protective eusociality found in termite colonies towards a more independent and self-sufficient lifestyle. However, there are a limited number of sequenced cockroach representatives, so further cataloging of diverse Blattodea genomes is necessary to pinpoint exact relationships between these gene families and cockroach-termite evolution. Overall, synteny and ortholog comparison between these Blattodea reveal possible mechanisms of divergence between termites and cockroaches and highlight the potential applications of a chromosomally resolved *P. americana* genome.

## 5 Summary

We sequenced the genome of the American cockroach using high fidelity PacBio long reads in conjunction with Hi-C Illumina short reads. This 3.23 Gb assembly is highly contiguous, with a contig N50 of 42 Mb and a scaffold N50 of 188 Mb, and 98.93% of the assembly is contained within 17 putative chromosomes. The quality of this assembly is further exemplified by its genomic and protein BUSCO scores, which are 99.7% and 97.8% complete respectively. This high-quality assembly, generated with cutting edge sequencing technology, is a substantial improvement over existing *P. americana* genomes, and we report an entire chromosome that was missing from a previously published assembly. This genome is expected to facilitate future study of cockroach physiology and Blattodea evolution.

## 6 Data Availability

Data associated with this study are available from the NCBI Sequence Read Archive under BioProjects PRJNA1098420 (principal haplotype) and PRJNA1098419 (alternative haplotype). Raw sequencing reads for Hi-C data is available from the SRA with accession SRX24490912, and PacBio HiFi reads may be obtained from accessions SRX24490909, SRX24490910, and SRX24490911. RNA-seq data is deposited under the SRA BioProject PRJNA1105088. Scripts used for assembling and analyzing this genome are available at: https://github.com/rldockman/PAMFEO.

## 8 Acknowledgements

We would like to thank Jeremy Schrader for his assistance with preparing and sequencing the DNA for this project.

## 10 Supplemental Tables and Figures

**Table S1:**
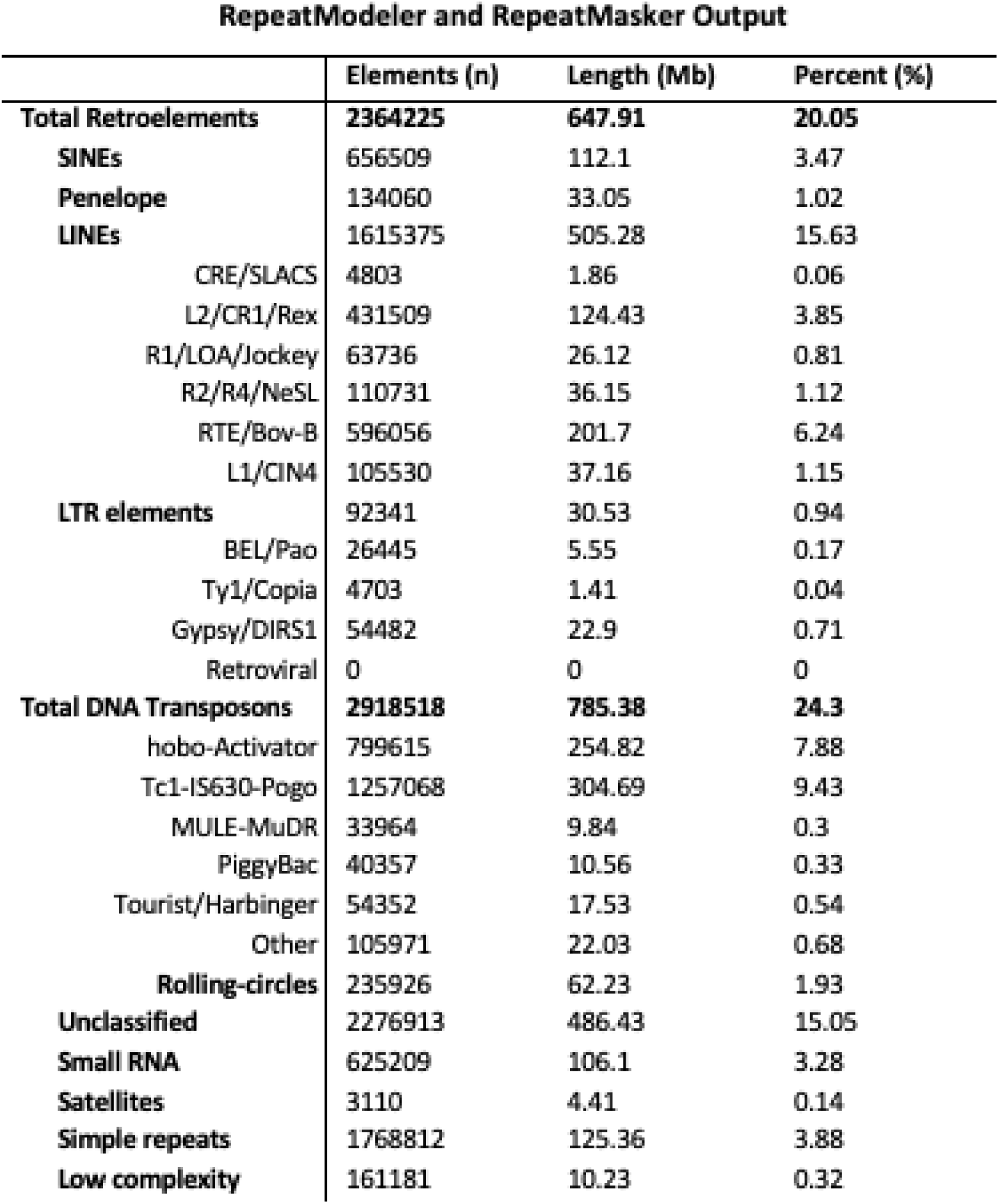
Summary of repeat classifications in *P. americana* that were identified with RepeatModeler and RepeatMasker.

**Figure S1:**
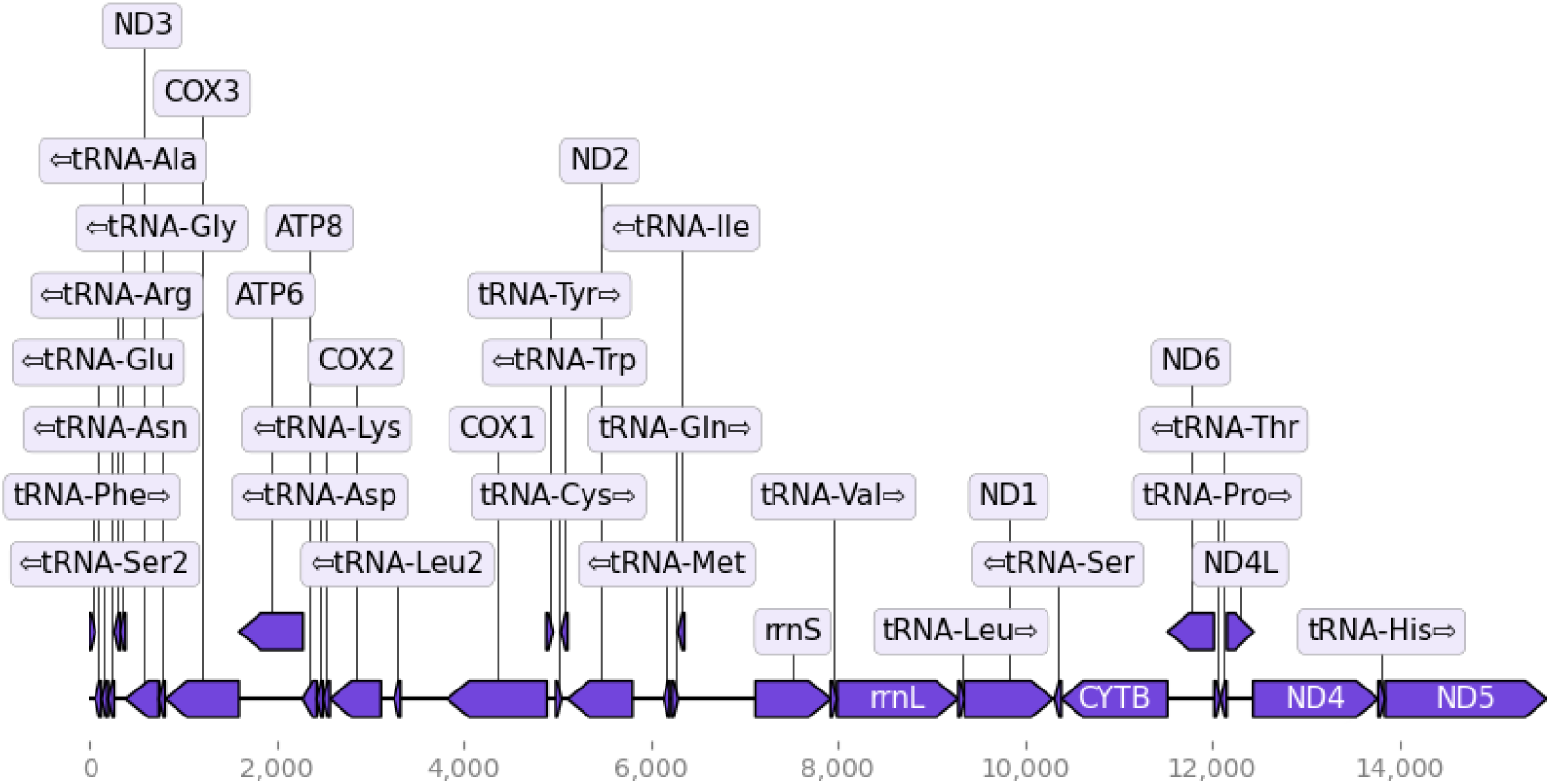
Mitogenome annotation output from MitoHiFi.

**Figure S2:**
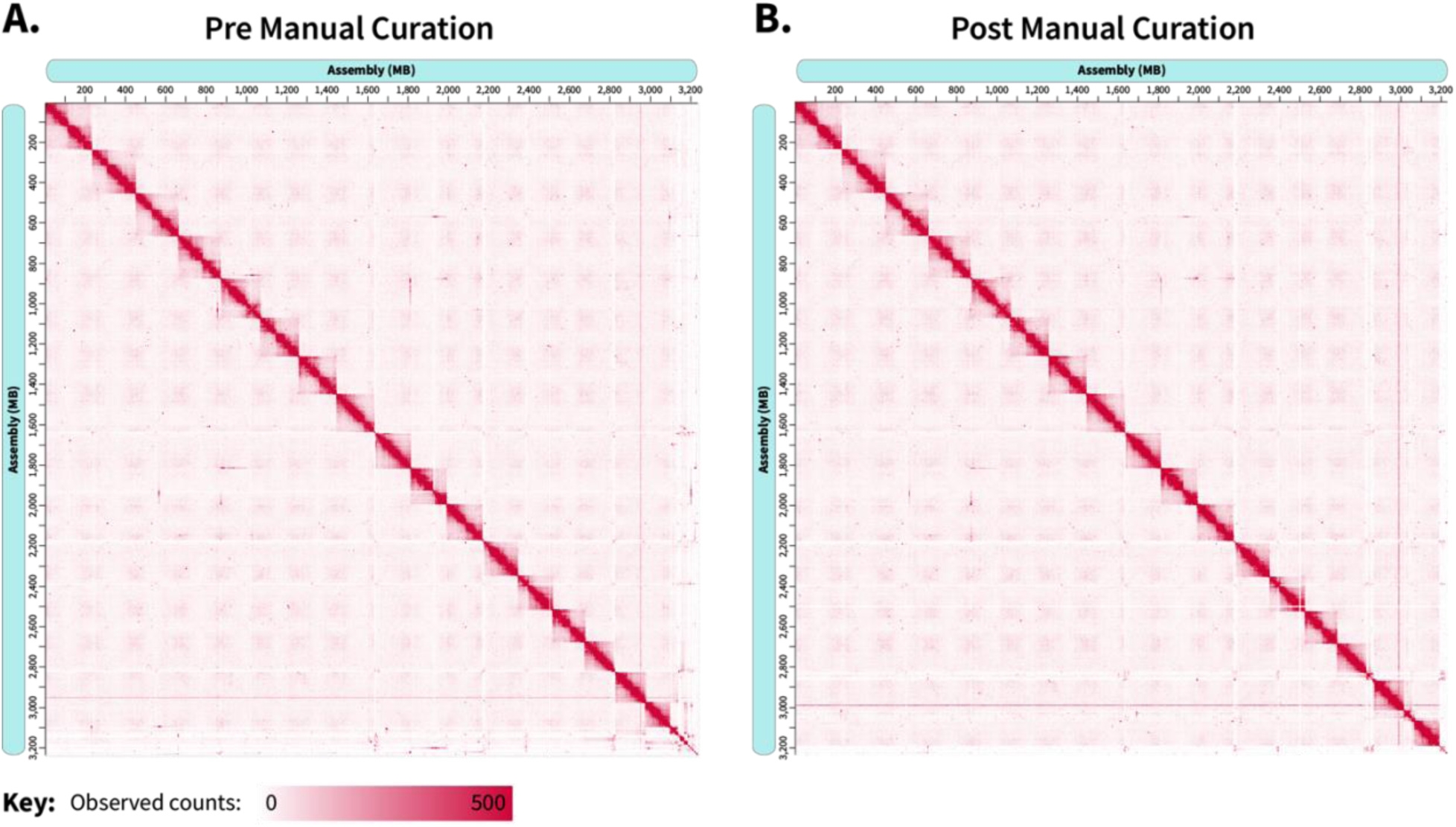
Hi-C contact maps of the assembled genome **(A)** before and **(B)** after manual curation using Juicebox.

**Figure S3:**
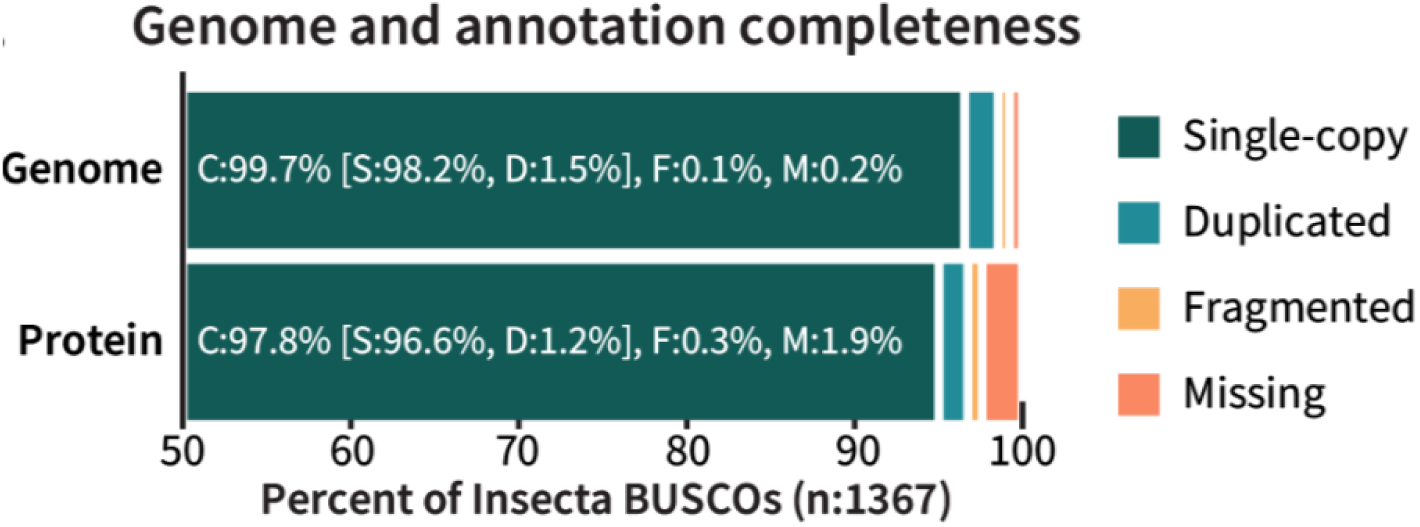
Insecta BUSCOs (version 10) identified in this *P. americana* genome assembly and protein annotation.

**Figure S4:**
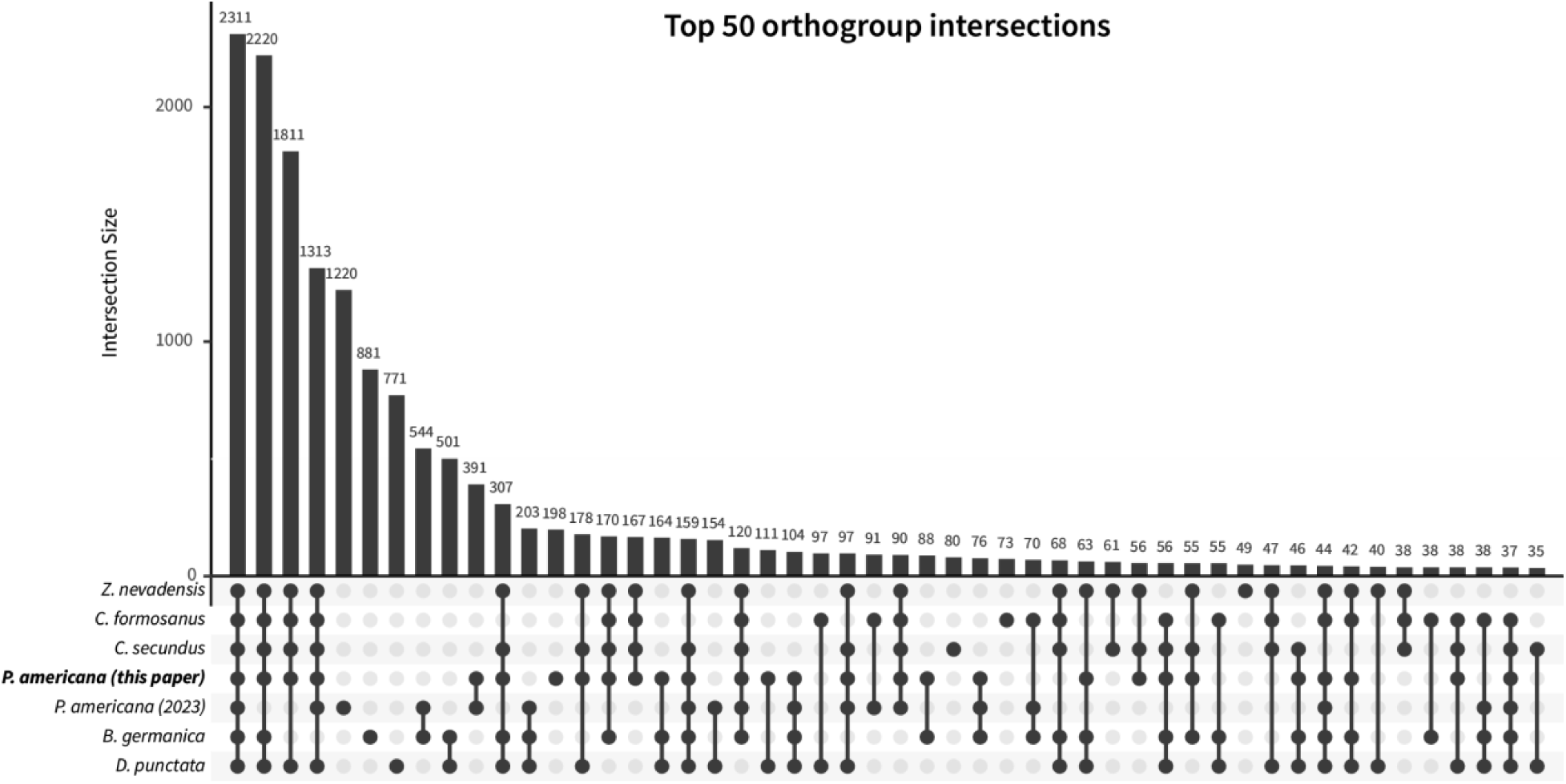
UpSet plot of orthogroup overlap between Blattodea species when the previous *P. americana* assembly is included.

**Figure S5:**
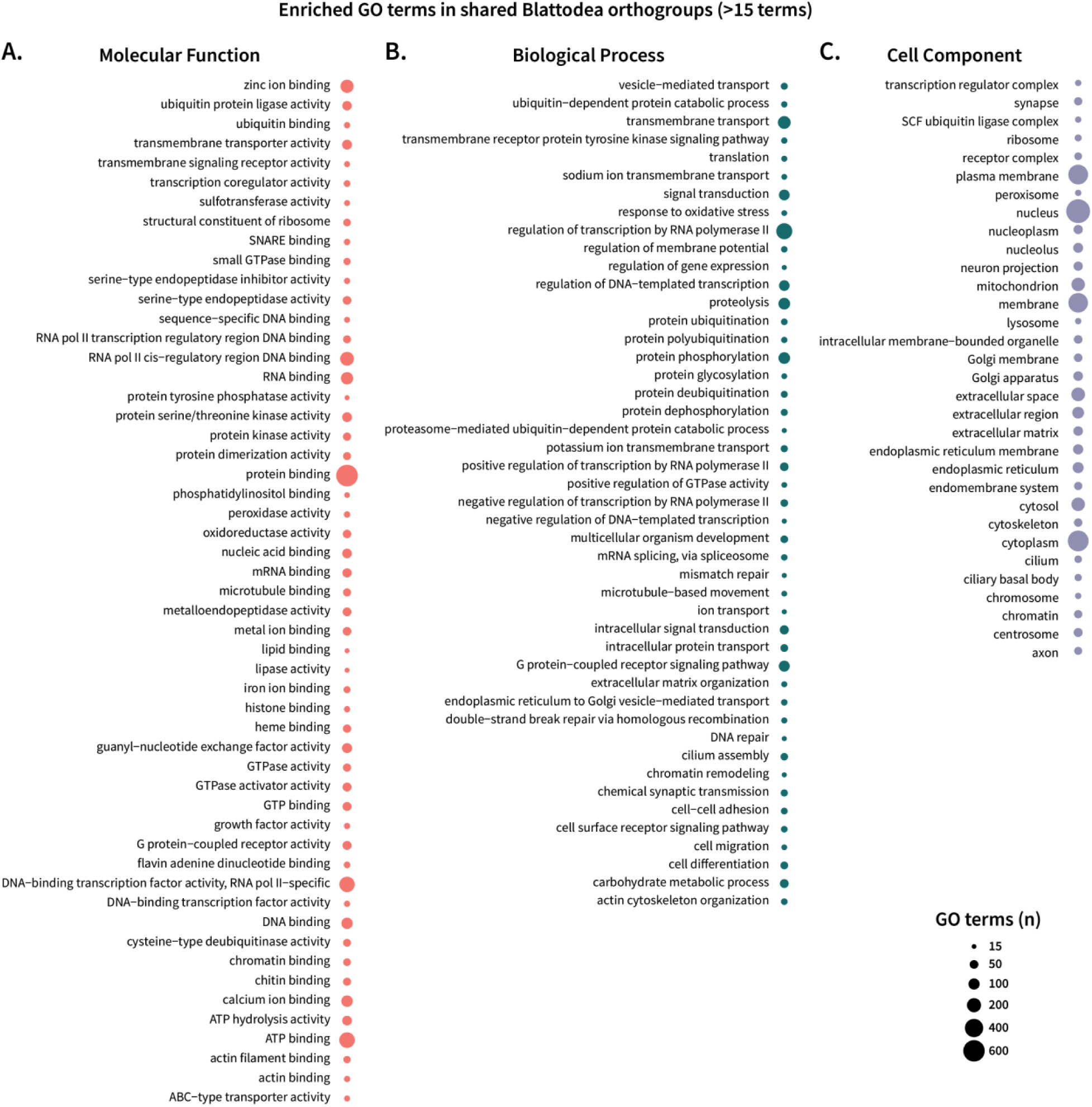
GO terms enriched in orthogroups shared by all Blattodea species. Genes were included in this analysis if they were sorted into the set of shared orthogroups in Figure 5D and had associated GO assignments. The profiles of enriched GO terms per category were generated in R with the package clusterProfiler. GO terms with abundances of at least 15 genes are reported for **(A)** molecular functions, **(B)** biological processes, and **(C)** cellular components.

**Figure S6:**
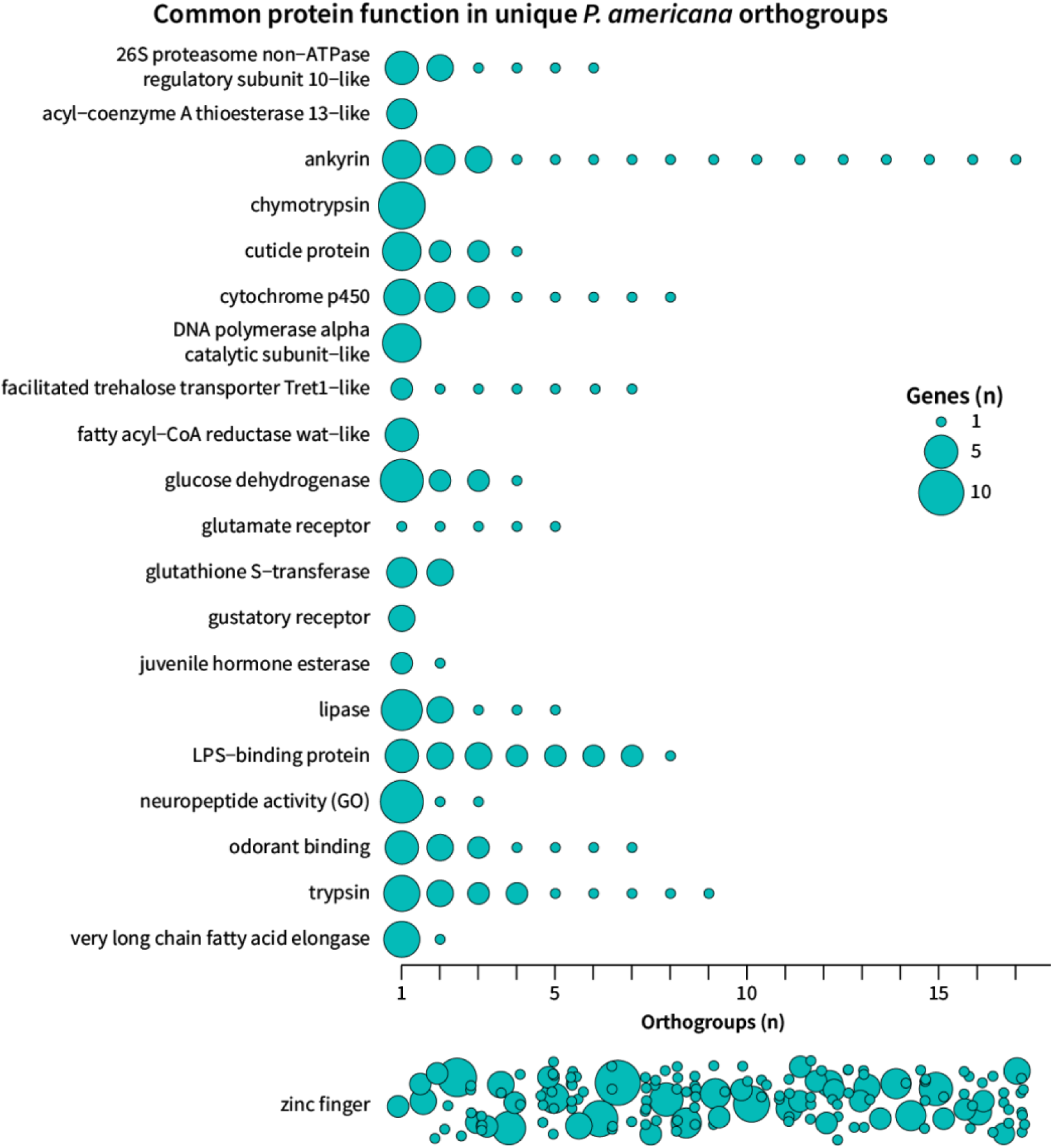
The names or GO terms of gene loci belonging to the 589 *P. americana*-unique orthogroups were collapsed to identify common functions in these genes and visualized with a bubble plot. Each bubble indicates a separate orthogroup per protein/function and is scaled by the number of genes sharing the associated name/GO term within the orthogroup. Proteins characterized by zinc finger motifs are plotted separately due to their large number of orthogroups (157 orthogroups, 248 genes).

